# Two Similarity Metrics for Medical Subject Headings (MeSH): An Aid to Biomedical Text Mining and Author Name Disambiguation

**DOI:** 10.1101/039008

**Authors:** Neil R. Smalheiser, Gary Bonifield

## Abstract

In the present paper, we have created and characterized several similarity metrics for relating any two Medical Subject Headings (MeSH terms) to each other. The article-based metric measures the tendency of two MeSH terms to appear in the MEDLINE record of the same article. The author-based metric measures the tendency of two MeSH terms to appear in the body of articles written by the same individual (using the 2009 Author-ity author name disambiguation dataset as a gold standard). The two metrics are only modestly correlated with each other (r = 0.50), indicating that they capture different aspects of term usage. The article-based metric provides a measure of semantic relatedness, and MeSH term pairs that co-occur more often than expected by chance may reflect relations between the two terms. In contrast, the author metric is indicative of how individuals practice science, and may have value for author name disambiguation and studies of scientific discovery. We have calculated article metrics for all MeSH terms appearing in at least 25 articles in MEDLINE (as of 2014) and author metrics for MeSH terms published as of 2009. The dataset is freely available for download and can be queried at http://arrowsmith.psych.uic.edu/arrowsmith_uic/mesh_pair_metrics.html.

## Background

Text mining analyses often involve estimating the similarity of two terms or concepts. In the biomedical domain, MEDLINE records include manual indexing by experts of topics discussed in each article, using a standardized hierarchical terminology of Medical Subject Headings (MeSH terms) that is employed to assist in retrieval of articles on a given topic. Various schemes have been proposed for relating different MeSH terms to each other in terms of their similarity. In general, these schemes can be classified as a) semantic, e.g., the path distance separating the two MeSH terms on the hierarchical tree; b) contextual, e.g., to what extent the two MeSH terms co-occur within the same articles; and c) lexical, e.g., the edit distance involved in transforming one term into another (Zhou et al, 2015). Co-occurring MeSH terms have been studied as an indicator of relations discussed in articles (e.g., Burgun and Bodenreider, 2001; Srinivasan and Hristovski, 2004; Kastrin et al, 2014) and MeSH-based similarity metrics have been employed in clustering of topically related articles (e.g., Lee et al, 2006; Zhu et al, 2009; Boyack et al, 2011). Several text mining models devoted to literature-based discovery have utilized similarity of two MeSH terms, or of two UMLS concepts, as features (e.g., Cohen et al, 2010; Theodosiou et al., 2011; Workman et al., 2013, 2015).

In the present work, we have computed and characterized two different MeSH term pair similarity metrics. The first involves calculating how often two different MeSH terms co-occur in the same articles, relative to the expected chance level (i.e., due to the frequencies of each MeSH term considered independently). We confirm that this metric captures topical similarity as judged by human raters, and point out some potential new uses for the metric in text mining. The second metric is novel: how often two different MeSH terms co-occur in the body of articles written by the same individual, relative to the expected chance level. As we will show, this author-based metric has potential value for author name disambiguation modeling. Both person-centered and article-centered metrics are being released openly as comprehensive datasets and can be viewed via public web interfaces at http://arrowsmith.psych.uic.edu/mesh_pair_metrics.html.

## Methods

### Article-based metric

For each article included in the 2014 baseline version of MEDLINE. we extracted the Medical Subject Headings (MeSH) associated with the article, and calculated the number of times that each pair of MeSH terms co-occurred within the same article, as well as the total number of articles in which each MeSH term occurred. A stoplist of the 20 most frequent MeSH terms (D’Souza and Smalheiser, 2014) was employed to remove them from consideration, since highly frequent terms would appear to be similar to all other MeSH terms. Only those MeSH terms appearing in at least 25 articles were considered in calculating term similarity measures, since lower values would be highly subject to noise. The final number of included MeSH terms is 25,548.

### Author-based metric

The 2009 Author-ity dataset (Torvik et al, 2005; Torvik and Smalheiser, 2009) is based on a snapshot of PubMed (which includes both MEDLINE and PubMed-not-MEDLINE records) taken in July 2009, including a total of 19,011,985 Article records, 61,658,514 author name instances and 20,074 unique journal names. Each instance of an author name is uniquely represented by the PMID and the position on the paper (e.g., 10786286 3 is the third author name on PMID 10786286). Thus, each predicted author-individual cluster is associated with a list of predicted PMIDs written by that individual. For each author-individual cluster included in 2009 version of Author-ity, we extracted the MeSH terms found in each cluster (each term is counted once in each cluster, regardless of how many articles it appeared in within that cluster). We then calculated the number of times that each pair of MeSH terms co-occurred within the same author-individual cluster, across all clusters in the dataset. Only MeSH terms that were included for calculating article-based similarity (see above) were considered for calculating author-based similarity; a total of 25,007 MeSH terms were included in the author-based metric.

There are 37,385,852 pairs of MeSH terms included in the article similarity metric. 201,136,960 pairs of MeSH terms were included in the author similarity metric. The number of pairs calculated for author metric is greater than included in the article metric, since MeSH terms were counted as co-occurring if they were mentioned in ANY articles written by a given individual, even if they never co-occurred in the same article. Conversely, the article metric contains 729,894 pairs that are not included in the author-based metric (i.e., involving MeSH terms which were added to MEDLINE after 2009). Finally, 36,655,958 pairs of MeSH terms were included in both Author and Article similarity metrics, and could be directly compared to see how the two metrics capture different aspects of similarity.

### Calculation of odds ratios

For any pair of MeSH terms, the number of co-occurrences needs to be normalized by the total number of occurrences of each MeSH term, in order to assess properly how meaningful it is to find two terms co-occurring (in the same article, or in the set of articles published by a given author). Two very common MeSH terms might be expected to co-occur often just by chance, whereas it will be highly significant if one observes any co-occurrence of two very rarely occurring MeSH terms. We computed the co-occurrence score that would be expected simply by chance (for two MeSH terms of their size), separately for the article-based and author-based metrics. This was done by ranking all MeSH pairs by the geomean of their individual document occurrences, dividing into bins of 5,000 pairs (i.e., each having roughly the same size), and calculating the average co-occurrence score across all MeSH pairs in the same bin. Finally, we calculated the MeSH odds ratio for each pair of MeSH terms present in that bin, by taking the observed co-occurrence score divided by the average co-occurrence score for that bin. This is similar to the manner in which odds ratios were computed for journal similarity metrics in D’Souza and Smalheiser (2014).

## Statistics

We employed correlation measures to characterize the relationship between two metrics, which allowed us to estimate the similarity of the metrics. In general, the nonparametric Spearman rho rank correlation coefficient is more appropriate for these comparisons, because the metrics are generally not linear. However, we also present the parametric Pearson r correlation coefficient as well, since there is some value in comparing the Pearson and Spearman values (e.g., if both are high, the relationships are likely to be linear, whereas if Pearson is very low and Spearman is very high, the relationships are likely to be nonlinear). Because each correlation was computed across millions of data points, statistical significance is generally extremely high and p-values are not displayed.

## Results

As one might expect, the article-based and author-based MeSH odds ratios were significantly correlated, but perhaps surprisingly, the correlations were only about 0.5 (Pearson r = 0.501, Spearman rho = 0.558). In other words, the two metrics do not simply measure the same thing. Rather, the tendency of two MeSH terms to co-occur in the same article reflects somewhat different aspects of similarity than the tendency of the same MeSH terms to co-occur within the body of work published by the same author.

The article-based metric, which counts co-occurrence of two MeSH terms in the same article, is subject to some limitations and constraints since a single article tends to have only 8-20 MeSH terms, and since MEDLINE indexers follow complex rules by which they decide to pick a given MeSH term (e.g., if more than one term is applicable but they lie vertically within the hierarchical tree, they are instructed to choose only the most specific term). Nevertheless, the article-based metric corresponded well to human ratings of semantic relatedness. Pedersen et al (2007) compiled a list of 29 UMLS concept (CUI) pairs annotated by physicians on a 1 to 4 scale of semantic similarity (Table 1). We mapped these to the corresponding MeSH term pairs as far as possible, and found that physician ratings correlated very well with the article-based metric (r = 0.67). A similar finding was observed with ratings by medical coders (Table 1).

**Table 1.**
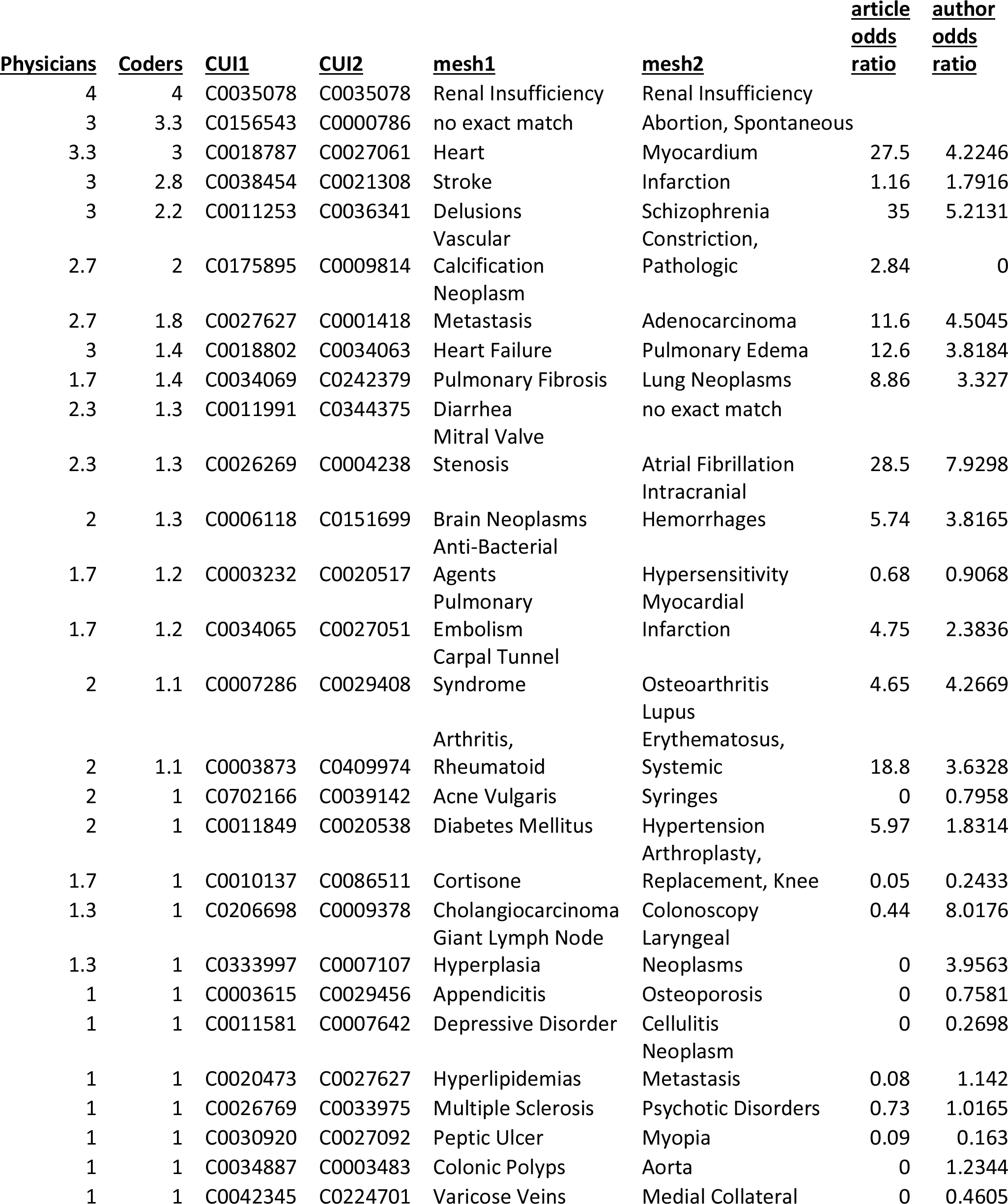
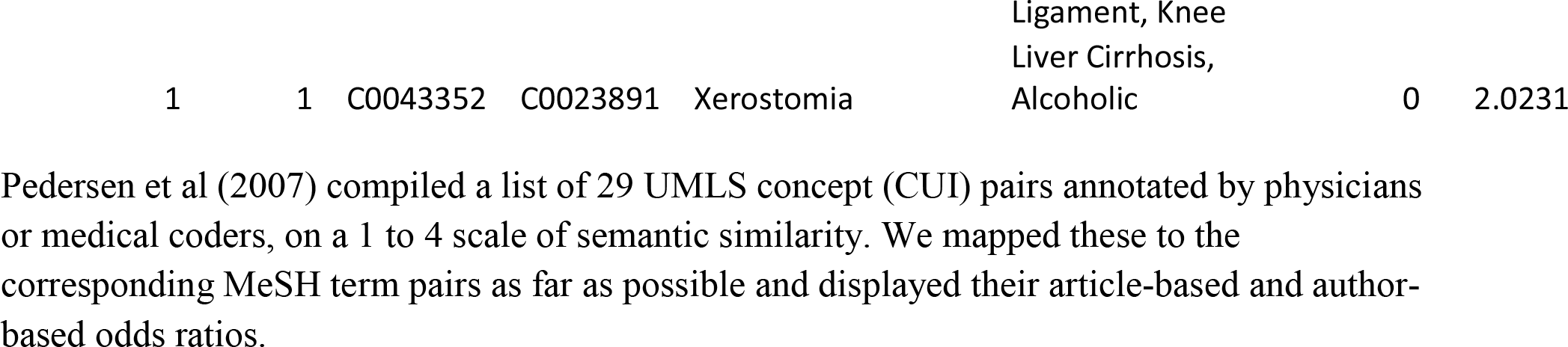
Comparison of article-based and author-based MeSH term pair odds ratios against human raters’ judgments of semantic relatedness.

In contrast, these ratings showed a much lower correlation with the author-based metric (r = 0.38). Note that one of the test pairs (Cholangiocarcinoma and Colonoscopy) co-occurred relatively infrequently within the same article (odds ratio = 0.44), but had a high author-based odds ratio (= 8.02), indicating that certain individuals, presumably GI specialists, tended to publish on both topics. Seven of the 29 MeSH pairs had no co-occurrences at all within the same article (and hence have article-based similarity scores of 0), yet all of these had author-based cooccurrences such that the odds ratios were greater than zero. This may suggest that the author-based metric is more sensitive in detecting indirect similarities.

Another feature of the author-based metric is its “smoothing” effect relative to the article-based metric. If an author has published 7 articles, and each has 8 MeSH terms, potentially there is a pool of 56 MeSH terms to be considered pairwise, compared to only 8 MeSH terms for each article. This makes the author metric relatively robust and less influenced by fluctuations due to low sampling, particularly for MeSH terms that occur in relatively few articles. For any given MeSH term, its article-based odds ratio tended to achieve higher maximal values than did the author-based odds ratios (article-based maximal odds ratio = 73.525 mean + 52.16 SD vs. author-based maximal odds ratio = 49.213 mean + 36.11 SD, a difference that is highly significant (p< 0.0001, one-tailed unpaired t-test)).

One way to compare the article-based and author-based metrics is to examine the datasets as they can be queried on the Arrowsmith project MeSH Pair Metrics page http://arrowsmith.psych.uic.edu/arrowsmith_uic/mesh_pair_metrics.html. The user selects any MeSH term from a drop-down menu, and the site displays the top 20 most related MeSH terms ranked according to either the article-based or author-based metrics (Table 2). For each MeSH pair, the site also displays the number of articles in which each MeSH term occurs, the number of co-occurrences (in articles or author-individual clusters), the average number of cooccurrences expected for two MeSH terms by chance (based on their size), and the calculated odds ratio. It is interesting to view how the article-based and author-based metrics sometimes emphasized different dimensions of similarity. For example, consider the top 20 terms related to the MeSH term “Tennis” (Table 3). The article-based metric lists 8 terms related to physical therapy and disorders that affect tennis players (vs. 4 terms listed under the author-based metric), whereas the author-based metric listed 10 other sports (vs. 5 sports listed under the article metric). Simply put, articles on tennis talked more about disorders afflicting tennis players, and did not generally include other sports in the same articles, whereas authors who wrote about tennis wrote more often about a variety of other sports. Another interesting example is “Abbreviations as Topic” (Table 4). The top 20 terms according to the article-based metric included 7 terms that were related to nursing and medications (vs. 2 listed under the author-based metric), whereas the author-based metric included 14 terms related to information science (vs. 8 under the article-based metric).

**Table 2.** The top 20 MeSH terms most similar to ‘Clergy’ [MeSH], ranked by article-based odds ratio.

| *Rank* | *MeSH Term* | *Article Count* | *Article Co-Occurrence* | *Article Odds Ratio* | *Author Co-Occurrence* | *Author Odds Ratio* |
| --- | --- | --- | --- | --- | --- | --- |
| 1 | Catholicism | 7383 | 403 | 73.9721 | 489 | 11.9222 |
| 2 | Pastoral Care | 3105 | 426 | 73.0453 | 541 | 28.6092 |
| 3 | Chaplaincy Service, Hospital | 924 | 180 | 68.9655 | 230 | 30.6748 |
| 4 | Religion and Medicine | 9795 | 216 | 43.1310 | 419 | 9.7469 |
| 5 | Spirituality | 4549 | 121 | 31.1054 | 188 | 9.2904 |
| 6 | Religion and Psychology | 5439 | 122 | 27.4899 | 288 | 10.0404 |
| 7 | Christianity | 6287 | 170 | 26.7632 | 353 | 10.4056 |
| 8 | Religion | 11340 | 156 | 25.9395 | 376 | 6.2134 |
| 9 | Protestantism | 693 | 50 | 20.6782 | 79 | 14.5488 |
| 10 | Professional Role | 7739 | 90 | 20.3712 | 150 | 4.2735 |
| 11 | Theology | 1141 | 49 | 19.0661 | 78 | 10.2821 |
| 12 | Child Abuse, Sexual | 8060 | 76 | 17.5115 | 112 | 3.0210 |
| 13 | Judaism | 2255 | 45 | 17.4014 | 133 | 9.7722 |
| 14 | Counseling | 26810 | 108 | 16.6821 | 308 | 3.6347 |
| 15 | Value of Life | 5338 | 56 | 15.6863 | 137 | 5.3399 |
| 16 | Anecdotes as Topic | 4470 | 46 | 14.4201 | 67 | 3.4561 |
| 17 | Terminally Ill | 5155 | 44 | 12.7536 | 136 | 5.3030 |
| 18 | Attitude | 37996 | 112 | 12.2351 | 397 | 2.7826 |
| 19 | Euthanasia, Passive | 5808 | 57 | 12.1899 | 142 | 4.7142 |
| 20 | Ethics | 9353 | 51 | 12.0796 | 161 | 3.9591 |

Shown are the top 20 MeSH terms that co-occurred with “Clergy” [MeSH], article count = 1641, ranked by article odds ratio. Author-based co-occurrences and odds ratios are also shown.

**Table 3.**
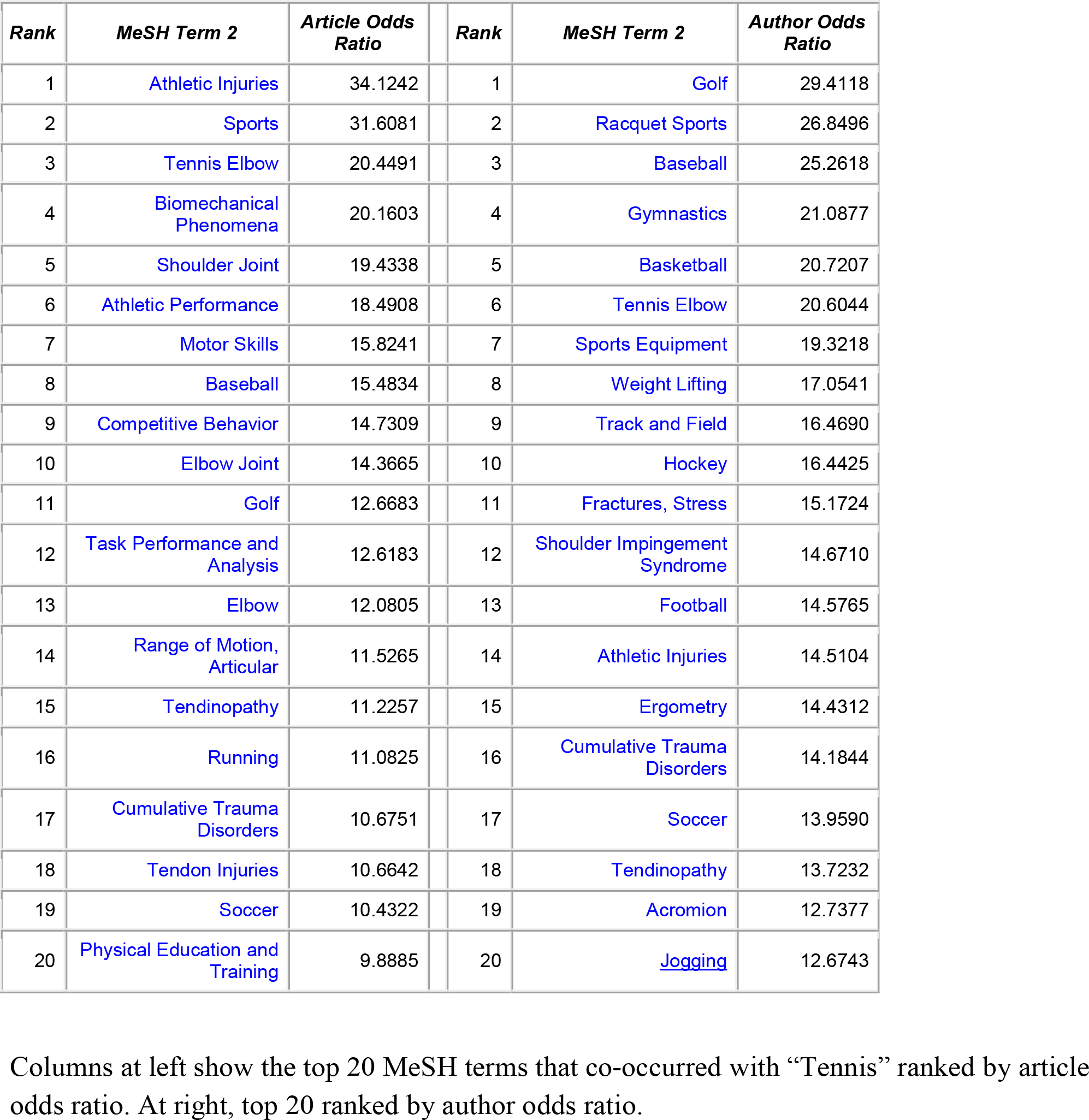
Top 20 MeSH terms most related to ‘Tennis’ [MeSH] by article and by author odds ratios.

| *Rank* | *MeSH Term 2* | *Article Odds Ratio* | *Rank* | *MeSH Term 2* | *Author Odds Ratio* |
| --- | --- | --- | --- | --- | --- |
| 1 | Athletic Injuries | 34.1242 | 1 | Golf | 29.4118 |
| 2 | Sports | 31.6081 | 2 | Racquet Sports | 26.8496 |
| 3 | Tennis Elbow | 20.4491 | 3 | Baseball | 25.2618 |
| 4 | Biomechanical Phenomena | 20.1603 | 4 | Gymnastics | 21.0877 |
| 5 | Shoulder Joint | 19.4338 | 5 | Basketball | 20.7207 |
| 6 | Athletic Performance | 18.4908 | 6 | Tennis Elbow | 20.6044 |
| 7 | Motor Skills | 15.8241 | 7 | Sports Equipment | 19.3218 |
| 8 | Baseball | 15.4834 | 8 | Weight Lifting | 17.0541 |
| 9 | Competitive Behavior | 14.7309 | 9 | Track and Field | 16.4690 |
| 10 | Elbow Joint | 14.3665 | 10 | Hockey | 16.4425 |
| 11 | Golf | 12.6683 | 11 | Fractures, Stress | 15.1724 |
| 12 | Task Performance and Analysis | 12.6183 | 12 | Shoulder Impingement Syndrome | 14.6710 |
| 13 | Elbow | 12.0805 | 13 | Football | 14.5765 |
| 14 | Range of Motion, Articular | 11.5265 | 14 | Athletic Injuries | 14.5104 |
| 15 | Tendinopathy | 11.2257 | 15 | Ergometry | 14.4312 |
| 16 | Running | 11.0825 | 16 | Cumulative Trauma Disorders | 14.1844 |
| 17 | Cumulative Trauma Disorders | 10.6751 | 17 | Soccer | 13.9590 |
| 18 | Tendon Injuries | 10.6642 | 18 | Tendinopathy | 13.7232 |
| 19 | Soccer | 10.4322 | 19 | Acromion | 12.7377 |
| 20 | Physical Education and Training | 9.8885 | 20 | Jogging | 12.6743 |

Columns at left show the top 20 MeSH terms that co-occurred with “Tennis” ranked by article odds ratio. At right, top 20 ranked by author odds ratio.

**Table 4.** Top 20 MeSH terms most related to ‘Abbreviations as Topic’ [MeSH] by article and by author odds ratios.

| *Rank* | *MeSH Term 2* | *Article Odds Ratio* |  | *Rank* | *MeSH Term 2* | *Author Odds Ratio* |
| --- | --- | --- | --- | --- | --- | --- |
| 1 | Terminology as Topic | 28.2360 |  | 1 | Dictionaries as Topic | 8.8602 |
| 2 | Medication Errors | 14.5379 |  | 2 | MEDLINE | 8.3068 |
| 3 | Nursing Assessment | 11.9482 |  | 3 | Subject Headings | 7.5937 |
| 4 | Periodicals as Topic | 10.9649 |  | 4 | Weights and Measures | 7.4758 |
| 5 | Drug Prescriptions | 10.2136 |  | 5 | Medical Subject Headings | 7.1207 |
| 6 | Weights and Measures | 9.1896 |  | 6 | Natural Language Processing | 6.8958 |
| 7 | Writing | 9.1089 |  | 7 | Metric System | 6.7002 |
| 8 | Medical Records | 7.7367 |  | 8 | Abstracting and Indexing as Topic | 6.6596 |
| 9 | MEDLINE | 7.4530 |  | 9 | Unified Medical Language System | 6.1576 |
| 10 | Language | 7.2093 |  | 10 | Databases, Bibliographic | 6.0713 |
| 11 | Communication | 6.9483 |  | 11 | International System of Units | 5.9347 |
| 12 | Joint Commission on Accreditation of Healthcare Organizations | 6.8768 |  | 12 | Dictionaries, Medical | 5.8901 |
| 13 | Natural Language Processing | 6.3139 |  | 13 | Names | 5.2069 |
| 14 | Abstracting and Indexing as Topic | 6.1107 |  | 14 | Wit and Humor as Topic | 4.5648 |
| 15 | Nursing Records | 5.9940 |  | 15 | Vocabulary, Controlled | 4.2828 |
| 16 | Safety Management | 5.8633 |  | 16 | Hypermedia | 4.2093 |
| 17 | Publishing | 5.6080 |  | 17 | Reminder Systems | 4.0059 |
| 18 | Names | 5.2402 |  | 18 | Patient Identification Systems | 3.8644 |
| 19 | Information Storage and Retrieval | 5.1120 |  | 19 | Peer Review, Research | 3.8040 |
| 20 | Handwriting | 5.0601 |  | 20 | National Library of Medicine (U.S.) | 3.6824 |

Columns at left show the top 20 MeSH terms that co-occurred with “Abbreviations as Topic” ranked by article odds ratio. At right, top 20 ranked by author odds ratio.

## Discussion

The present paper describes and provides comprehensive article-based and author-based similarity metrics for pairs of Medical Subject Heading (MeSH) terms. We also present a web query interface that allows users to retrieve, for any specified MeSH term, the top 20 most related MeSH terms according to either the article-based or author-based metric.

As discussed in the introduction, article-based MeSH term pair similarity has been previously discussed by others and utilized in studies of information retrieval, document clustering, and literature-based discovery. Here, we have confirmed that the article-based metric does correspond to judgments of semantic relatedness made by human raters. We note that indexing an article with two MeSH terms suggests that the article may discuss a potential relation between the two topics - especially if the article-based odds ratio of that pair is greater than 1, i.e., if the two MeSH terms co-occur in the same articles more often than expected by chance. We deem a MeSH term pair as “important” if their article odds ratio >1. This further suggests a new kind of similarity metric for relating different articles to each other. That is, for any pair of articles, one can score the number of “important” MeSH term pairs that they share, perhaps weighted so that MeSH term pairs with higher odds ratios count more. Counting MeSH term pairs will be more stringent and restrictive than counting individual shared MeSH terms.

The author-based MeSH term pair similarity measures the tendency of the same individual to discuss two different topics at some time during their career, i.e., in the body of articles that they have authored or co-authored. This says as much (or more) about the individual, and his or her range of interests, as it does any overt relation between the two topics. For example, the first author of this paper (NS) has written on a variety of subjects ranging from extracellular matrix biochemistry to natural language processing to a biography of the early neuroscientist Walter Pitts. The two MeSH terms “Dystroglycans” and “History, 20th Century” are not obviously related to each other, and in fact, do not co-occur in any single article in PubMed. Yet one might hypothesize that an investigator who has worked on both topics might be psychologically or otherwise better poised to detect new knowledge that bridges these two fields, or that requires assembling different pieces of knowledge from each field, than someone who has only worked in one field. Although novel discoveries often involve combining topics together in new ways (e.g., Uzzi et al, 2013; Chen, 2014; Mishra and Torvik, 2014), one can hypothesize that MeSH term pairs which do not co-occur at all in the same articles, yet have high author-based odds ratios, may draw upon a pool of prepared minds (to quote Pasteur) and be particularly likely to be linked in new discoveries. Those which have very low author-based odds ratios might be less likely to be assembled into a new finding by an individual investigator. Multi-disciplinary teams may have a particular advantage for the investigation and discovery of findings that involve putting together topics that have very low author-based odds ratios. These hypotheses are testable, and may further our understanding of why and how particular novel combinations of topics lead to new discoveries.

Our original reason for studying author-based MeSH term similarity was to create an additional feature that can be used to disambiguate author names on PubMed articles comprehensively (Torvik et al., 2005; Torvik and Smalheiser, 2009). The most relevant measure of similarity for disambiguation is not topic centered, but rather author-centered. For example, two journals may cover the same topic (e.g., Scandinavian Journal of Immunology vs. Iranian Journal of Immunology), yet the same person may have very little likelihood of publishing articles in both journals. Thus, the 2009 Author-ity disambiguation dataset was earlier mined to create a metric that comprehensively measures the tendency of individuals to publish articles in any two journals (D’Souza and Smalheiser, 2014). Here, the author metric measures the tendency of individuals to publish a body of articles that is indexed by any two different MeSH terms during their careers.

We are currently using the author-based metric in modeling author disambiguation, as follows: Given two articles sharing the same author (lastname, firstinitial), we compute all MeSH terms for each article and consider the pairs that are formed across articles (i.e., one MeSH term in article 1 paired with another MeSH term in article 2). The three highest author-based odds ratios are averaged and used as the similarity score for the two articles. This feature is one of several new features that we are using to update and improve the performance of our author name disambiguation model.

## Implementation

The datasets and readme.pdf are freely available for download from the Arrowsmith project website (http://arrowsmith.psych.uic.edu/arrowsmith_uic/mesh_pair_metrics.html) as well as from the UIC Institutional Repository, INDIGO, under the terms of the Creative Commons Attribution-NonCommercial-ShareAlike CC BY-NC-SA license 4.0. The MeSH term pair data is contained in mesh_pair_metrics.txt (16 GB uncompressed, 5 GB compressed).

## Acknowledgements

This research was supported by National Institutes of Health grants R01LM010817 and P01AG039347. We thank Vetle I. Torvik for advice regarding computing, and Arushi Chawla for assistance in data analysis.

## Authors’ contributions

NRS designed the experiments. GB performed the data analyses and created the datasets. NRS wrote the paper.

